# On the Interaction of Biopotential Sensing and Right Leg Drive System with Electro-Quasistatic Human Body Communication

**DOI:** 10.1101/2022.06.13.495999

**Authors:** Shreeya Sriram, Kurian Polachan, Shreyas Sen

## Abstract

Continuous long-term sensing of biopotential signals is vital to facilitate accurate diagnosis. The current state of the art in wearable health monitoring relies on radiative technology for communication. Due to their radiative nature, these systems result in lossy and inefficient transmission, limiting the device’s life span. Human Body Communication has emerged as an energy-efficient secure communication modality, and literature has shown body communication to transmit biopotential signals at 100x lower power than traditional radiative technologies. Unlike radiative communication that uses airwaves, HBC, specifically Capacitive Electro-Quasistatic HBC (EQS-HBC), couple signals and confine them within the human body. In Capacitive EQS-HBC, the transmitter uses an electrode to modulate the body potential to transmit data. The modulation of body potential by HBC raises the following concerns. Will HBC transmissions affect the quality of biopotential signals sensed from the body? Additionally, since biopotential sensing systems commonly use Right Leg Drive (RLD) to bias body potential, there is also a concern if RLD can affect the quality of HBC transmissions.

For the first time, our work studies the interactions between EQS-HBC and biopotential sensing. Our work is important since understanding HBC-RLD interactions is integral to developing EQS-HBC-based biosensors for Body Area Networks (BANs). For the studies, we conducted lab experiments and developed circuit theoretic models to back the experimental outcomes. We show that due to their higher frequency content and common-mode nature, HBC transmissions do not affect the differential sensing of low-frequency biopotential signals. We show that biopotential sensing using RLD affects HBC. RLD deteriorates the signal strength of HBC transmissions. We thus propose not to use RLD with HBC. We demonstrate our proposed solution by transmitting ECG signals using HBC with 96% correlation compared to the traditional wireless system at a fraction of the power.

## I. Introduction

FUTURE connected healthcare applications like remote health monitoring demand the design of wearable devices for long-term and continuous monitoring of biopotential signals such as Electrocardiogram (ECG) or neural activity. Wearable devices in these applications sense vital signs at periodic intervals and transmit these readings in real-time to a nearby hub or cloud for further processing and logging. Continuous monitoring of vital signs using wearable devices is especially beneficial to individuals with chronic conditions and allows for the timely detection of changes in health conditions. For instance, continuous monitoring can track changes in the patient’s health and detect any sudden changes that may occur either during sleep or during levels of heightened activity.

Current state of the art in wearable technology for biopotential sensing includes smart watches such as the Apple Watch, Fitbit Sense, Samsung Galaxy Watch to name a few. These devices are capable of recording the ECG signals by placing the fingertip on the crown or dial of the device allowing for an on-demand ECG signal. These devices record single lead, two electrode ECGs with one electrode below the watch surface, in contact with the skin surface and the the dial of the watch acting as the second electrode. Though popular among the public, such devices only record 30 s worth of ECG information and do not continuously monitor the patient’s ECG. Continuous monitoring devices such as holter monitors [1] or ECG patches on the other hand have a limited life span. For instance, the Vital Patch, recently introduced by VitalConnect which continuously monitors Electrocardiogram (ECG), has a limited life span of only six days. The reason for the limited life span of these wearables can be attributed to their use of power-hungry radiative electromagnetic (EM) communication technologies (e.g., WiFi, BLE) for the continous transmission of sensed biopotential data. In comparison to the sensing and processing modules in these wearables, the radiative communication module typically consumes 2-to-3 order of magnitude more power, making it the biggest bottleneck in achieving long term operation of the wearable. The radiative communication technologies consumes significant power due to the need for up-conversion of the baseband signal to higher frequencies and due to the loss associated with radiated airwaves.

Human Body Communication (HBC) which uses the conductive tissues of the body to transmit signals has emerged as an energy efficient alternate to radiative communication technologies for wearable designs requiring longer life span’s [2]–[4]. Unlike radiative communication that uses airwaves, HBC, specifically, Capacitive HBC operating in the electro quasistatic region (EQS-HBC), referred to as Capacitive-EQS-HBC uses the body as a wire to couple low frequency (<1 MHz) signals and contain them within the human body [5]. An on-body hub (e.g., a smartwatch) with an HBC receiver can read these signals and relay the information to a cloud for further processing. Recent studies have demonstrated Capacitive-EQS-HBC with sub-*µ*W of power consumption, 100x lower than BLE [4], [6]. Capacitive-EQS-HBC also provides improved physical security to sensitive health data by containing the communication signals within the body, thereby eliminating potential eavesdropping from nearby attackers [5].

Capacitive-EQS-HBC uses an electrode to couple signals to the body [7]. The transmitter uses this electrode to modulate the body potential to communicate data bits. For example, the transmitter may apply a higher voltage to the electrode and raise the body potential to convey bit 1 and apply a lower voltage to the electrode and lower the body potential to convey bit 0. The receiver uses an electrode to sample these body potential variations to infer the transmitted data. Here, modulating body potential by Capacitive-EQS-HBC for transmitting data raises the following important question — Can HBC transmissions affect sensing of body’s biopotential signals. Further, since HBC transmissions are dependent on the body’s potential, and since many of the biopotential sensing circuits biases the body potential by means of Right-Leg Drive (RLD) circuitry, it is also a question to ponder if biasing of body potential by RLD can impact HBC transmissions.

For the first time, our work investigates the effect of Capacitive-EQS-HBC on biopotential signal sensing and the effect of RLD of biopotential sensing circuits on Capacitive-EQS-HBC. Due to their higher frequency content, we show that Capacitive-EQS-HBC transmissions do not affect the sensing of low-frequency biopotential surface signals such as ECG. We show that RLD affects Capacitive-EQS-HBC. RLD deteriorates the signal strength of Capacitive-EQS-HBC transmissions. We thus propose not to use RLD with Capacitive-EQS-HBC and substitute its functionality with software filters. We argue that our proposed solution is a viable option for biopotential sensing systems with wearable form factors. Further, we demonstrate the solution by sensing ECG signals without RLD and transmitting the ECG data using Capacitive-EQS-HBC with 96% correlation compared to the traditional wireless system at a fraction of the power.

### A. Key Contributions

Following are the key contributions of this paper —

- Conclusively establishes that Capacitive-EQS-HBC does not interfere with biopotential signal measurements from the body.
- Analyses the effect of the Right Leg Drive (RLD) on Capacitive-EQS-HBC and justify its elimination for biopotential signal measurements in the presence of Capacitive-EQS-HBC.
- Demonstrate a proof-of-concept wearable that senses biopotential signals without RLD using filters to reduce noise in the absence of RLD. Experimentally verify Capacitive-EQS-HBC as a communication modality for biopotential signal transmission by comparing its quality with BLE.

### B. Outline

We organize this paper as follows: In Section I-C, we list related works on HBC focusing on biopotential sensing applications and how they differ from this paper’s work. Section II, provide background information on Human Body Communication (HBC) and Right-Leg Drive (RLD) for biopotential sensing. Section III discusses the need to investigate the interactions between Capacitive-EQS-HBC and biopotential sensing in a Body Area Network (BAN). In this section, we also analyze the effect of Capacitive-EQS-HBC on biopotential sensing and the effect of RLD on Capacitive-EQS-HBC. We conclude that Capacitive-EQS-HBC does not interfere with biopotential sensing, but RLD does interfere with Capacitive-EQS-HBC increasing channel loss. As a solution, in Section IV, we propose the elimination of RLD for biopotential sensing in wearables supporting Capacitive-EQSHBC. Further, In Section V, we develop a proof-of-concept wearable to demonstrate our proposed solution. We conclude our work in Section VI.

### C. Related Work

Several works in the literature demonstrate the use of HBC to transmit data around the body [8], [9]. However, only a few demonstrate the use of HBC in biomedical nodes that sense and transmit biopotential signals. In nodes that perform biopotential sensing and HBC, sensing and communication can interact and corrupt signal measurements and data.

Wang et al. [10] described a technique for biopotential sensing and HBC. In this technique, ECG signals acquired from the subject were transmitted through the body using impulse radio On-Off Keying. Intelligent switching between the sensing and transmitting electrodes were employed to sense and transmit the signals. However, the work uses the galvanic mode of HBC, which uses a pair of electrodes to communicate data, limiting HBC transmissions’ range. Authors in [11] also demonstrate time-multiplexed sensing, and HBC transmissions. In this technique, the nodes perform biopotential signal measurements for a specific duration. Following the sensing cycle, the node transmits biopotential signals through the body. The method ensures that sensing and communication interact at no instance of time.

The time-multiplexed approach of sensing and HBC transmissions can avoid possible interaction between sensing and communication at the node level. However, the method cannot avoid the possibility of sensing and communication interactions between nodes when multiple sensor nodes are present on the body as in an HBC enabled BAN. Further, time-multiplexed sensing and communication are not practical for healthcare applications requiring continuous monitoring of biopotential signals; nodes need to stop signal measurements while performing communication. Tang et al. [12] describe a method for simultaneous body communication and EEG sensing on a node; however, they do not study the effect of body communication on biopotential sensing and vice-versa.

## II. Background: HBC and RLD

### A. Human Body Communication

HBC uses the body’s conductivity as a low-loss broadband channel to transmit signals. Electro-Quasistatic Human Body Communication (EQS-HBC), operates at frequencies below 1MHz, allowing for ultra-low power body communication at 414nW [6] and energy consumption of 6.3pJ/bit [13]. In the past, HBC at lower frequencies was unfavourable due to higher channel losses caused by the resistive 50 Ω terminations used at the receiver-side [14]. Maity et al. demonstrated high-quality HBC at low frequencies by replacing traditional 50 Ω terminations with capacitive termination. Capacitive termination reduces channel loss in the low frequency region by 100x [5], [13], [15] and thereby allowing ultra-low-power communication across the body. EQS-HBC operates in two different modes: Capacitive and Galvanic as shown in Figure 2.

**Fig. 1.**
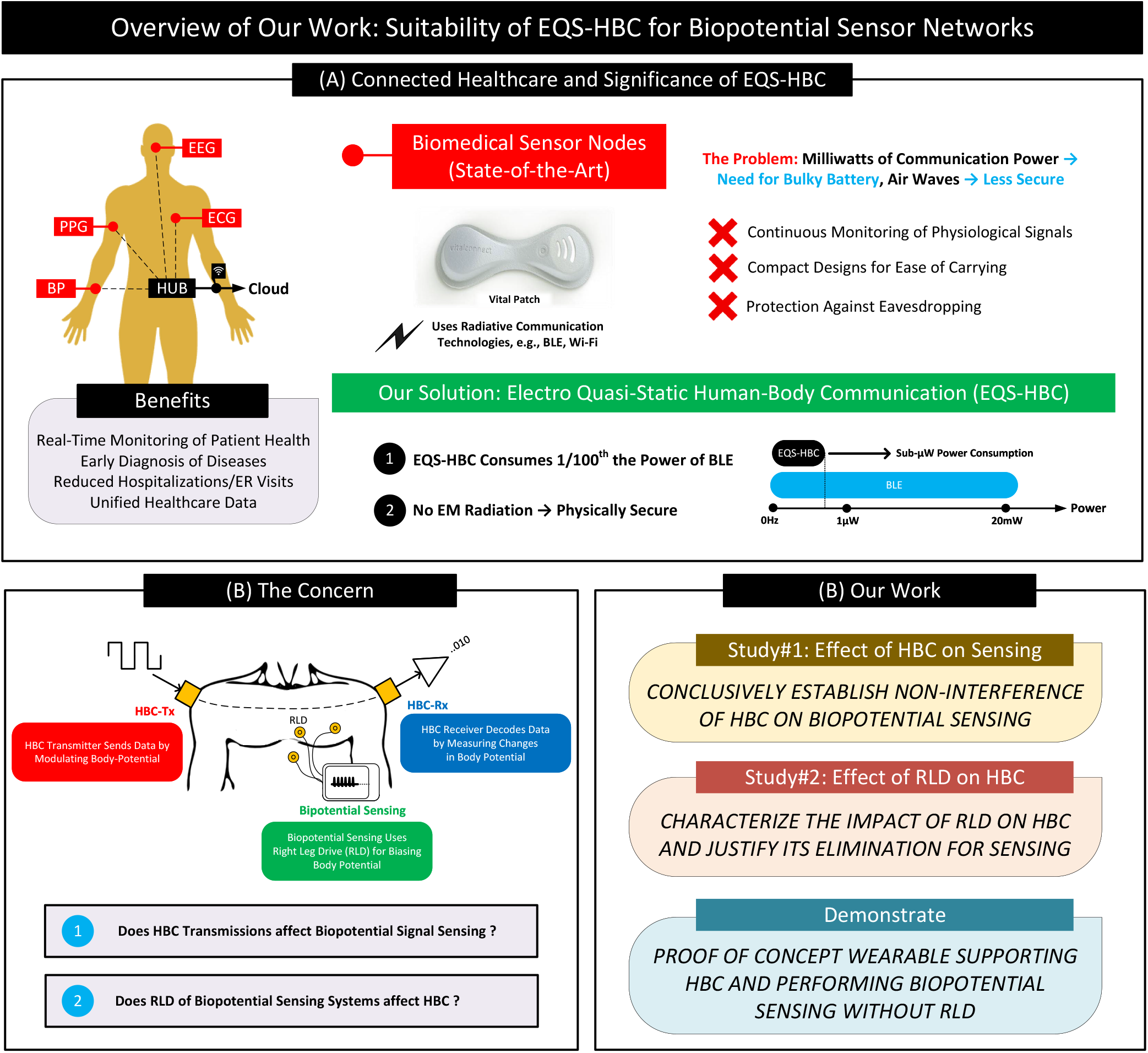
(A) State-of-the-Art biomedical sensor nodes use radiative communication technologies consuming milliwatts of power requiring bulky batteries for operation. They are also less secure due to the radiation of EM waves. As a solution, we propose EQS-HBC. (B) Capacitive-EQS-HBC and Right Leg Drive (RLD) of biopotential sensing systems modulate body potential, causing undue interactions between Capacitive-EQS-HBC and sensing. (C) Overview of our work — Our work studies the effect of Capacitive-EQS-HBC on sensing and RLD on Capacitive-EQS-HBC.

**Fig. 2.**
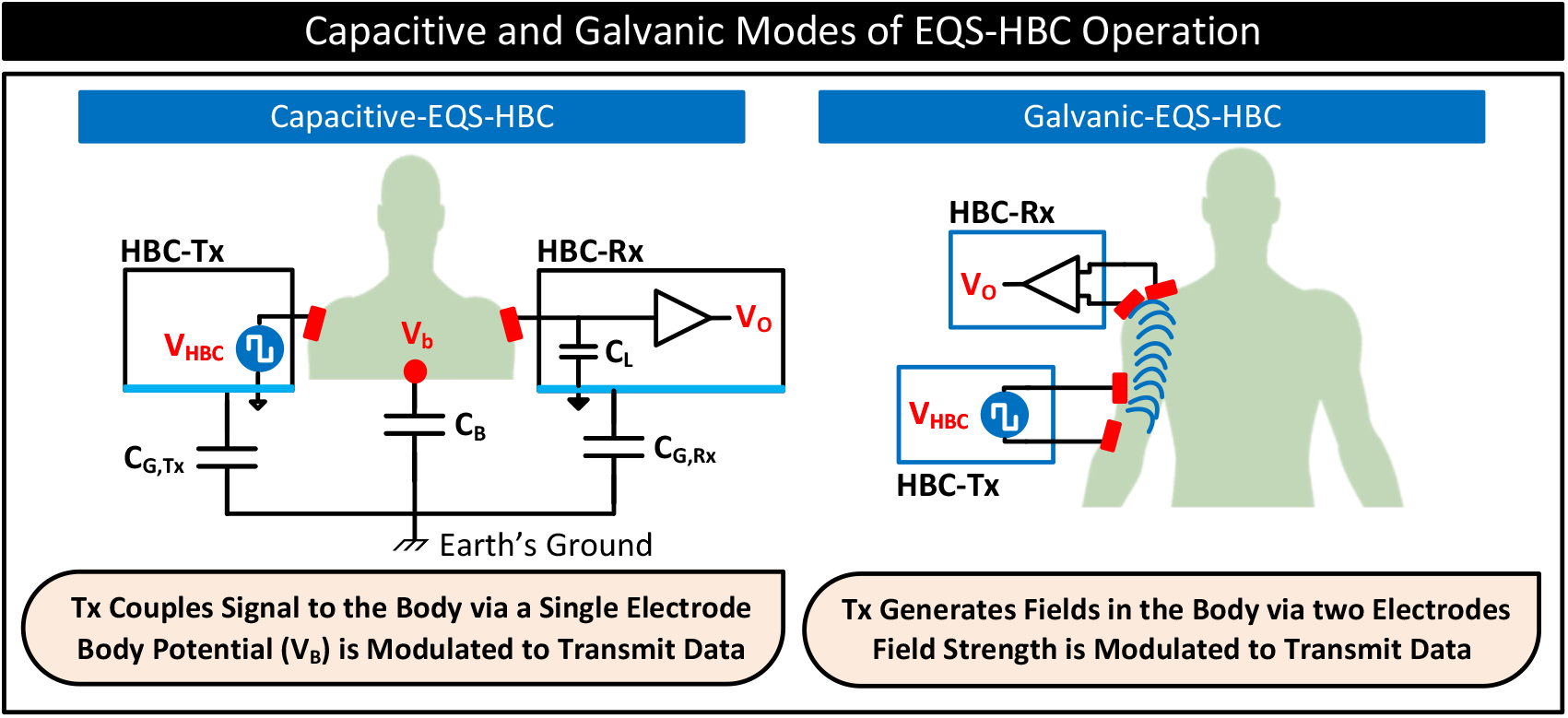
Different types of EQS-HBC: EQS-Capacitive-HBC, which transmits data by modulating body potential, and EQS-Galvanic-HBC, which transmits data by generating an electric field in the body and modulating its strength.

#### Capacitive-EQS-HBC

In capacitive-EQS-HBC, the transmitter uses a single electrode to couple signals to the body. The transmitter uses this electrode to modulate the body potential, (*V*_*B*_), to communicate data bits: 0 and 1. A receiver on the body detects these changes in the body potential to infer the data. The channel gain depends on the transmitter and receiver ground-to-earth capacitances (*C*_*G,T x*_, *C*_*G,Rx*_), body capacitance, *C*_*B*_, and receiver-side termination, *C*_*L*_ as follows:

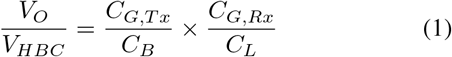

#### Galvanic-EQS-HBC

In contrast to the capacitive mode of EQS-HBC operation discussed above, EQS-HBC can also operate in galvanic mode. In Galvanic-EQS-HBC, the transmitter uses a pair of electrodes to couple signals to the body. The transmitter uses these electrodes to generate fields inside the body and modulate their strengths to communicate data bits. The configuration results in electric field lines concentrating near the proximity of the transmitting node, reducing the signal strength to deteriorate with distance from the transmitter, limiting the transmission range [16]–[18].

Our work focuses on the capacitive mode of EQS-HBC and its interaction with biopotential sensing. ***In the remainder of this paper, unless explicitly specified, when referring to HBC, we refer to the capacitive-EQS-HBC***.

### B. RLD for Biopotential Sensing

Biopotential sensing uses electrodes in contact with the body to sense signals from within the body to infer bodily activities [19]–[21]. Figure 3 (A) shows a simplified scheme to measure electrocardiogram (ECG), a biopotential signal representing heart activity [22]. The scheme uses two electrodes, *E*1 and *E*2, attached to the surface of the chest and separated by a distance to measure ECG. Differencing the signal voltages, *V*_1_ and *V*_2_ from the electrodes *E*1 and *E*2 yield the characteristic electrocardiogram (ECG) waveform. In measurement setups that use a difference amplifier, the common-mode voltage at the amplifier input terminals, *V*_*CM*_ = (*V*_1_ + *V*_2_)*/*2 = *V*_*b*_ *− V*_*m*_, plays a significant role. For accurate difference measurements, the *V*_*CM*_ has to be within the input common-mode range of the difference amplifier. For a rail-to-rail, single supply (*V*_*DD*_) difference amplifier, *V*_*cm*_ cannot be below *V*_*K*_ or above *V*_*DD*_ *− V*_*K*_; *V*_*K*_ being the amplitude of the ECG signal. Otherwise, the output of the difference amplifier will not faithfully track the differences in *V*_1_ and *V*_2_ and ECG. In biopotential measurement systems, the body’s potential defines *V*_*CM*_. Without any driving source connected to the body, noisy external sources, e.g., AC mains, define *V*_*CM*_. Parasitic capacitors *C*_*intf*_, *C*_*B*_, *C*_*L*_ and *C*_*G*_ in Figure 3 (A) fixes *V*_*CM*_ due to AC power line interference. Depending on their values, *V*_*CM*_ may go outside the required input common-mode range of the difference amplifier.

**Fig. 3.**
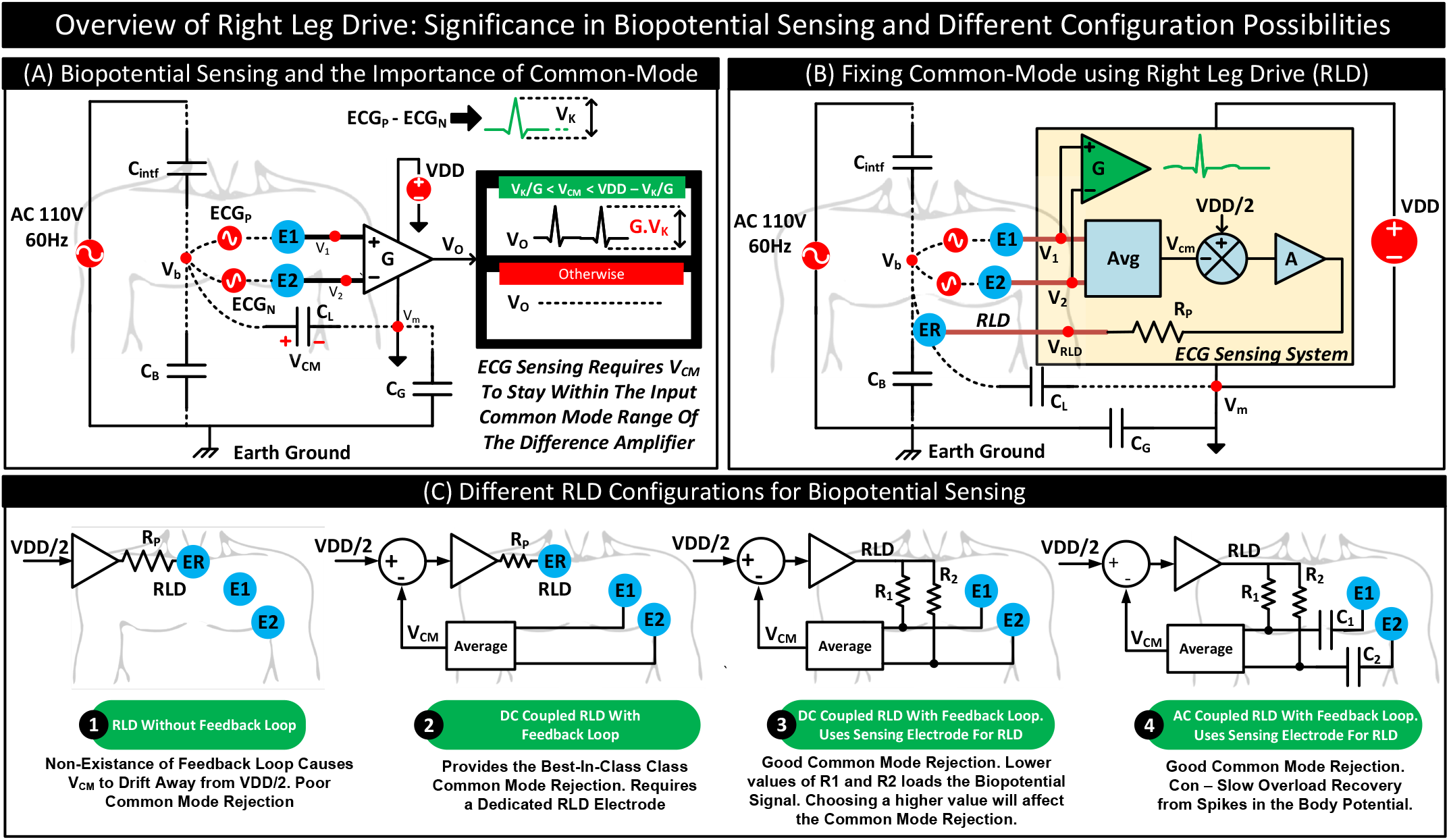
Power line interference model of an ECG sensing system performing a single lead ECG measurement. For accurate ECG measurements, *V*_*CM*_ has to be within the input common-mode range of the difference amplifier. (B) ECG sensing system with Right Leg Drive (RLD) to fix *V*_*CM*_ to 0.5*V DD*. The RLD here uses a dedicated electrode *ER* and a feedback loop to fix *V*_*CM*_ against variations in the body potential due to power line interferences. (C) Different RLD configurations reported in the literature.

As a solution, biopotential sensing systems use a Right leg drive (RLD) to fix *V*_*CM*_ [20], [23], [24]. The electrode, ER, attached to the body in Figure 3 (B) represents the RLD electrode. The RLD electrode fixes the voltage across *C*_*L*_ i.e., *V*_*CM*_, to center around the input common-mode range of the difference amplifier, i.e., 0.5*V*_*DD*_. Typically, A PI controller-based closed-loop circuit drives the RLD electrode to fix *V*_*CM*_ to 0.5*V*_*DD*_. PI controller is necessary for two purposes. First, it helps accurately fix *V*_*CM*_ to 0.5*V*_*DD*_ using the RLD electrode. Second, due to finite impedances between *E*1, *E*2, and RLD electrode with the body, the RLD electrode voltage, *V*_*RLD*_, does not necessarily match *V*_*CM*_. The use of the PI controller ensures that the magnitude of *V*_*RLD*_ is adjusted to set *V*_*CM*_ to 0.5*V*_*DD*_. PI controller also helps nullify variations in the body’s potential, *V*_*b*_, with external noise sources such as AC mains. AC mains can cause variations in the *V*_*b*_ and thus *V*_*CM*_ at 60 Hz. *V*_*CM*_, in this case, may go out of the input common-mode range of the difference amplifier. Even if *V*_*CM*_ is within the input common-mode range, due to the limited CMRR of the difference amplifier (and the passives), variations in *V*_*CM*_ may propagate to the output and affect the measured ECG signal quality. The feedback loop using the PI controller ensures *V*_*CM*_ stay at any instance of time close to 0.5*V*_*DD*_. The feedback loop thus works to reduce the common mode variations as seen at the input of the difference amplifier. In short, RLD serves two purposes. First, it fix *V*_*CM*_ to the input common-mode range of the difference amplifier. Second, it nullify noise sources, particularly power line noises, varying *V*_*CM*_ and corrupting ECG due to the limited CMRR of the difference amplifier.

Figure 3 (C) shows different RLD configurations reported in the literature [25]. Figure 3 (C)-1 represents the case where the RLD uses the electrode *ER* to drive 0.5*V DD*. The method does not use a feedback loop, which results in poor common-mode rejection. Figure 3 (C)-2 represent the RLD configuration of Figure 3 (B). It uses *ER* and a feedback loop to fix *V*_*CM*_. The configuration offers the best-in-class common-mode rejection. Figure 3 (C)-3 represents the RLD configuration which does not use a dedicated RLD electrode. Instead, the RLD in this configuration uses ECG electrodes *E*1 and *E*2 to fix *V*_*CM*_. To avoid RLD from corrupting ECG, resistors *R*_1_ and *R*_2_ are used between the RLD driver and electrodes *E*1 and *E*2 respectively. Here, using lower values of *R*_1_ and *R*_2_ loads the ECG signals, and higher values result in lower common-mode rejection. The configuration, however, is useful in applications that want to avoid a dedicated RLD electrode. Figure 3 (C)-4 shows another configuration that does not require a dedicated RLD electrode. The configuration uses capacitors *C*_1_ and *C*_2_ to AC couple ECG signals to the difference amplifier. The AC coupled configuration offers good common-mode rejection. However, a spike in the body potential due to a pacemaker or a defibrillator can couple through *C*_1_ and *C*_2_ to the difference amplifier saturating its output. It will take long before the capacitor’s discharge, and ECG measurements resume.

## III. HBC *↔* Sensing Interactions

### A. The Need to Investigate

HBC modulates body potential to communicate data bits. On the other hand, biopotential sensing commonly uses RLD to fix body potential to a set value. When these communication and sensing occur simultaneously, body potential remains unpredictable. Fluctuations in body potential resulting from HBC can appear as common mode noise to the biopotential sensing systems resulting in noisy measurements. On the other hand, the feedback loop associated with RLD, which compensates for varying body potential, can nullify HBC signals deteriorating the quality of HBC transmissions. In short, simultaneous sensing and communication can affect both biopotential signal measurements and the quality of HBC.

A possible solution is to avoid these interferences by purposefully avoiding simultaneous sensing and communication. However, in many scenarios, such as in HBC enabled Body Area Networks (BAN), consisting of several on-body sensor nodes and an on-body hub supporting HBC, simultaneous sensing and communication are advantageous and sometimes unavoidable, motivating the need to study the interaction mechanisms between HBC and RLD and possible ways to minimize their effects. The following paragraphs list various possibilities for sensing and communication in HBC-enabled BAN, highlighting cases where simultaneous sensing and communication are advantageous.

#### a) Asynchronous Sensing and Communication

All nodes perform sensing and communication asynchronously. Here, the frequency modulation technique may be adopted to avoid interferences when two nodes communicate data simultaneously to the hub. HBC transmissions from each node to the hub may use distinct preassigned carrier frequencies. However, the method cannot avoid simultaneous sensing and communication scenarios and the resulting interactions. For instance, a node may perform biopotential sensing while another node transmits data to the hub and vice versa. Though it may be possible for nodes to detect the presence of communication signals and start sensing only when the channel is idle, the technique cannot guarantee the possibility of starting a communication from another node in the middle of sensing. The method also suffers from bandwidth issues. Since EQS-HBC operates at lower frequencies, the possibility of assigning distinct frequencies reduces the bandwidth that can be allowed for each node. The restricted bandwidth may limit the frequency or rate of data transfers depending on the number of nodes in the BAN. However, the method does not restrict the rate at which nodes perform sensing, which is advantageous when nodes perform intelligent analysis on the data locally and communicate only critical events to the hub.

#### b) Synchronous Sensing and Communication

Here, the hub may synchronize the sensing and communication of all the nodes. The hub may send an HBC beacon addressing specific nodes one at a time. Upon receiving the beacon, the nodes may perform sensing and then communicate data to the hub. A variation of this method is time-division multiplexing. The nodes may be time-synchronized, and each node uses a specific slot in a cycle for sensing and subsequent communication of data to the hub. In both cases, each node needs to wait for other nodes’ sensing and communication to complete, effectively reducing the sensing and communication rate proportional to the number of nodes in the BAN. Further, the synchronization requirement increases the hardware complexity of the nodes and the hub. Nevertheless, synchronous sensing and communication eliminate the possibility of simultaneous sensing and communication scenarios.

#### c) Asynchronous Sensing and Synchronous Communication

Here, the nodes perform sensing at any rate they consider necessary. However, the communication happens at specific instances of time. As in the previous case, the hub may initiate communication from nodes by sending beacons, or nodes may time-synchronize themselves and use a slot to transmit data to the hub. Though asynchronous sensing and synchronous communication may result in simultaneous sensing and communication, the method does not restrict the sampling rate; it only restricts the communication rate. In many cases, this is acceptable as nodes may not transmit each sample to the hub; instead, they may perform local processing and transmit only critical events to the hub.

#### d) Synchronous Sensing and Asynchronous Communication

In this method, nodes perform sensing at specific instances of time. The hub can either define the instances by sending beacons or by the nodes themselves if they are already time-synchronized. For instance, the hub can send a beacon addressing all the nodes to start sensing. The nodes, however, communicate asynchronously. Here, frequency modulation can help avoid collision of communication; however, the method cannot prevent the occurrence of simultaneous sensing and communication.

Table I summarizes the four possible modes of sensing and communication in an HBC-enabled BAN, their shortcomings, and the possibility of sensing/communication interactions. Note that three out of the four possible modes in the table results in simultaneous sensing and communications.

**Table I.** Different Possibilities for sensing and communication in a BAN.

| Possibilities |  | Rate Limitation |  | Sensing/Comm. Interaction |
| --- | --- | --- | --- | --- |
| Sensing | Comm. | Sensing | Comm. |  |
| ASYN | ASYN | × | ✓ | ✓ |
| SYNC | SYNC | ✓ | ✓ | × |
| ASYN | SYNC | × | ✓ | ✓ |
| SYNC | ASYN | ✓ | ✓ | ✓ |

Although the above discussions focused on the sensing and communication interaction between different nodes in a BAN, the simultaneous sensing/communication interactions also occur at the node level. When either sensing or communication happens asynchronously, a node can choose to perform the sensing and communications operations in two parallel processes (or threads), resulting in sensing and communication on the node to collide.

In short, there are two possibilities of sensing and communication interactions in a BAN. (i) A node performs sensing while another node performs HBC., and (ii) A node performs both sensing and HBC simultaneously. These possibilities lead to the following four different effects:

1. **Effect of a node’s HBC on another node’s sensing**.
2. Effect of a node’s HBC on the node’s sensing.
3. Effect of a node’s sensing on another node’s HBC.
4. **Effect of a node’s sensing on the node’s HBC**

In our work, we study the cases 1). and 4). Specifically, in 1), we study if HBC transmissions by a node can affect another node’s biopotential readings (i.e., ECG readings). In 4), we study if RLD used by a node for biopotential sensing affects its HBC transmission characteristics. We believe that the effect of a node’s HBC transmission on the node’s biopotential sensing can be minimized using a low-pass filter at the analog front-end for biopotential sensing. Due to their higher frequency content, the low pass filter may block HBC signals and avoid its interaction with sensing. Further, due to isolated grounds of on-body nodes, the effect of a node’s sensing on another node’s HBC is minimal. Nevertheless, to confirm our beliefs, we plan to investigate cases 2) and 3) in future works.

### B. Effect of a node’s HBC on another node’s Sensing

Since HBC transmissions modulate body potential, questions arise regarding whether HBC can affect the biopotential readings (e.g., ECG) in scenarios where HBC transmissions and sensing coincide. In this section, we conduct two experiments to understand the possible effects.

The first experiment uses a grounded medical-grade device similar to those used in hospitals to measure ECG in the presence of HBC transmissions. The objective of the experiment is to determine if HBC transmitters on a subject’s body can affect the measurement of biopotential signals in clinical settings such as in hospitals or ambulatory settings. Figure 4 (a) shows the experimental setup. It consists of a medical-grade ECG measurement device from MindRay technologies to measure single-lead ECG signals from a subject. On the subject, an on-body HBC transmitter continuously couples OOK modulated HBC signals to the body. We conclude from the ECG signals displayed on the measurement device (see Figure 4 (a)) that HBC do not affect grounded ECG measurements.

**Fig. 4.**
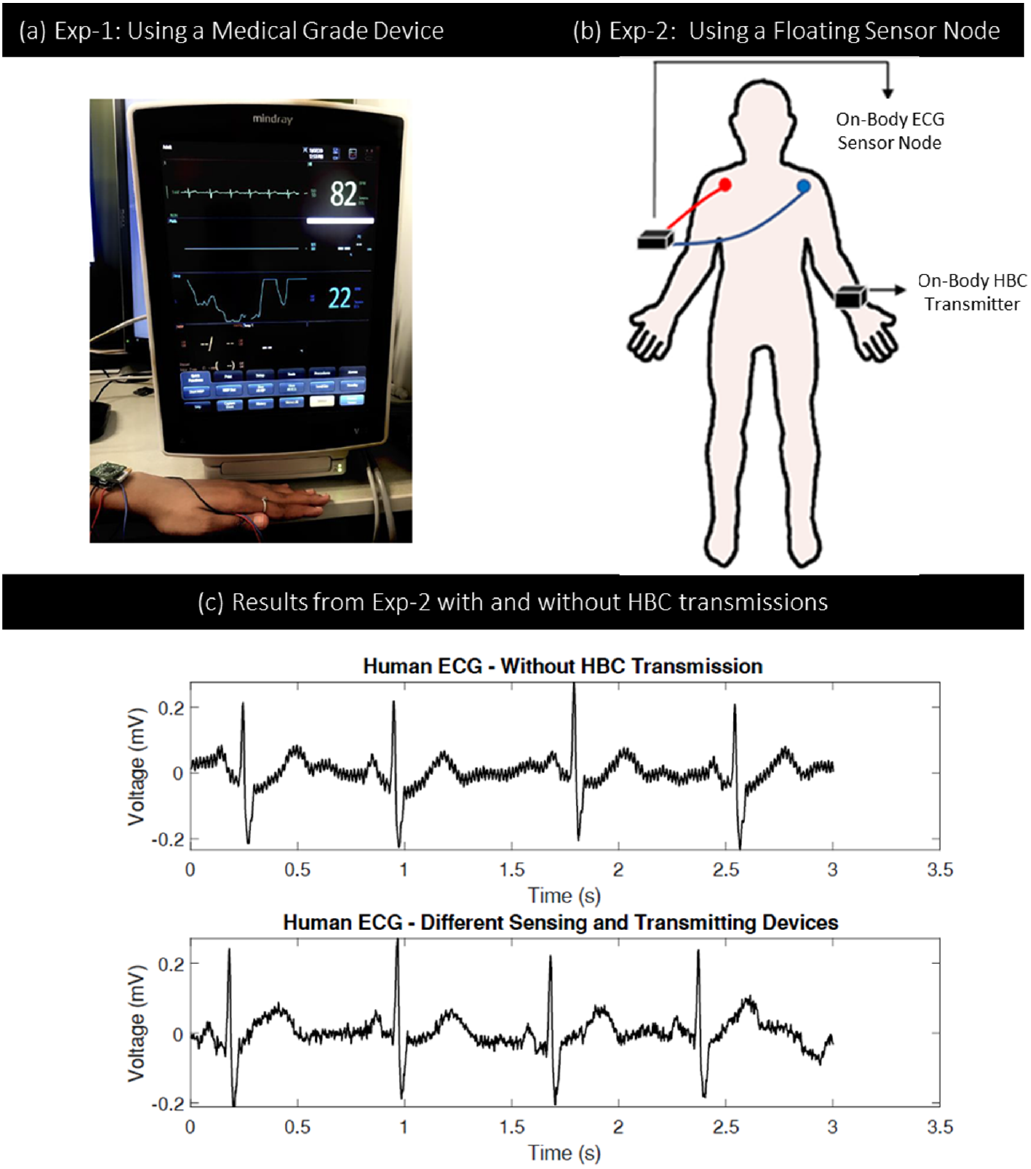
Effect of HBC on Biopotential Sensing: a) Experimental setup using a medical-grade device from MindRay technologies. The setup uses an on-body HBC transmitter to continuously couple OOK modulated HBC signals to the body. Clean ECG signals are observed in the device display. b) Experimental setup using a floating ECG sensor node consisting of an on-body HBC transmitter to continuously couples OOK modulated HBC signals to the body. c) ECG signals from the floating ECG sensor node remain clean with and without HBC transmissions.

The second experiment is a repetition of the first but uses a floating ECG sensor node (designed in-house) to substitute the grounded medical-grade device. The floating ECG sensor node mimics today’s popular wearable ECG sensor patches; the experiment thus helps understand the suitability of building HBC-based BANs comprising biopotential sensors of wearable form factor. Figure 4 (b) shows the experimental setup. It consists of a custom-designed ECG sensor node to measure single-lead ECG signals from the subject and an on-body HBC transmitter to continuously inject OOK modulated HBC signals to the subject’s body. Figure 4 (c) shows the results. We find that ECG signals are not affected by HBC.

#### a) Conclusion

Our findings based on the above experiments are as follows. Even if HBC transmissions collide with ECG measurements, they do not affect the ECG signal quality irrespective of whether the sensing system is grounded (as in a clinical setting) or floating (as in wearables). The reason can be attributed to the relatively high frequency of HBC transmissions (100 KHz - 1 MHz) compared to biopotential signals such as ECG (< 250 Hz), owing to which the sensing systems block HBC interferences using their front-end antialiasing filter. Although the experiments above study simultaneous sensing and communication interaction between two different nodes, we expect the results to remain the same, even when the same node carries out the sensing and communication operations. Here too, the antialiasing filter will help eliminate the interference from HBC transmissions on biopotential signal measurements.

### C. Effect of a node’s RLD on the node’s HBC

For the case of a node that performs simultaneous sensing and communication, the presence of the RLD will offer a low resistance return path to its HBC transmit signals, leading to higher HBC channel loss. This section derives the theory behind this behaviour and relates it with experimental findings.

#### 1) Theory

Figure 5 (A) shows the HBC-Tx side circuit model that performs simultaneous ECG sensing (with RLD) and HBC transmissions. In the model, *R*_*ER*_, *C*_*ER*_, *R*_*EH*_, and *C*_*EH*_ represent the resistances and capacitances between the RLD and HBC-Tx electrodes to the body, respectively. Here, *R*_*ER*_ and *R*_*EH*_ model the resistance of the electrode, the skin and the body tissue, while *C*_*ER*_ and *C*_*EH*_ model the capacitance of the electrode, and the skin [26]. *R*_*P*_ represents the protection resistor used to limit the current from the RLD driver passing through the patient’s body. Parameter *A* represents the gain of the RLD feedback loop. Since the feedback loop is designed to operate with limited bandwidth (<< *F*_*HBC*_), at the HBC frequency of *F*_*HBC*_, the impedance, *Z*_*p*_, seen by the electrode ER to *V*_*m*_ will remain *Rp* (and not *Rp/A*). *V*_*HBC*_ represents the HBC transmit signal. Capacitors 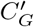 and *C*_*G*_ represent the parasitic capacitors between the positive and negative terminals of *V*_*HBC*_ to earth’s ground, respectively.

**Fig. 5.**
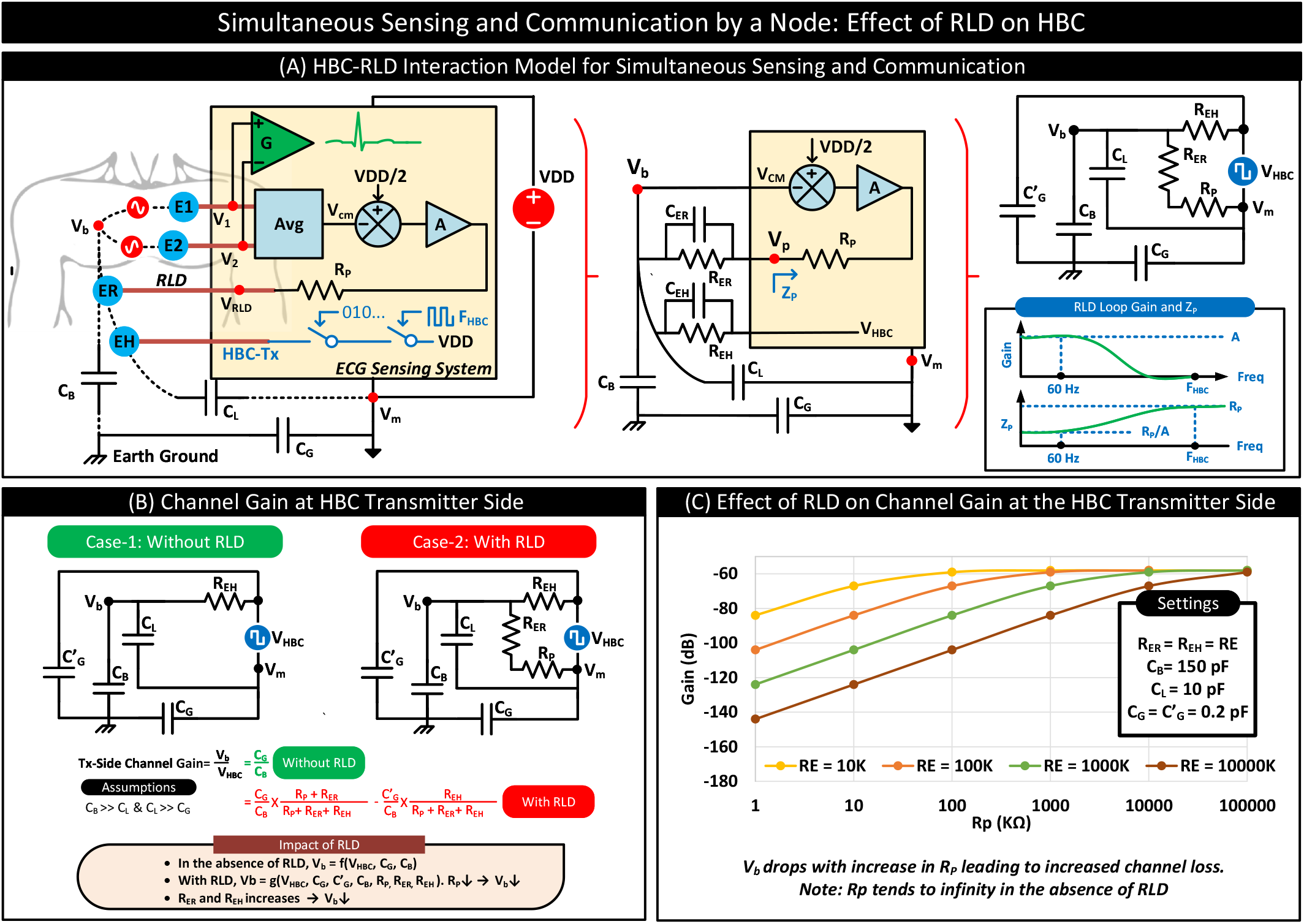
(A) HBC-RLD interaction of a node performing simultaneous ECG sensing and HBC transmissions. (B) Circuit model and channel gain equations with and without RLD (C) Plots showing the variation in channel gain with the RLD protection resistor, *R*_*p*_, and RLD/HBC electrode-to-body resistances, *R*_*ER*_ = *R*_*EH*_ = *RE*. Plots indicate that the RLD (*R*_*p*_ < *∞*) and *RE* affects the channel gain. Channel gain drops with lower *R*_*p*_ and higher *RE*.

Figure 5 (B) shows the HBC-Tx side circuit models with and without RLD. At low frequencies (< 1 MHz), the effect of *C*_*ER*_ and *C*_*EH*_ on HBC are insignificant; hence we remove these parasitics from the circuit model. In the absence of RLD, see Case-1 in Figure 5 (B), the HBC source drops its full potential, *V*_*HBC*_, across *C*_*L*_, which divides between *C*_*G*_ and 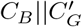. Since in Capacitive-HBC, the receiver decodes data by sensing *V*_*b*_, i.e., the voltage across *C*_*B*_, the resulting transmitter-side channel gain is 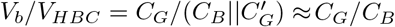 (Note: 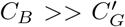). Note that in Case-1, the HBC source drops more or less the full potential across *C*_*L*_ even at frequencies above 1 MHz. Since, at higher frequencies, the voltage across *C*_*L*_ is dictated by *V*_*HBC*_*C*_*EH*_*/*(*C*_*L*_ +*C*_*EH*_) and *C*_*EH*_ is 100x higher than *C*_*L*_.

In the presence of RLD, see Case-2 in Figure 5 (B), RLD attached to the body creates a low-resistance return path for the transmitted HBC signals through the body tissues, mimicking a pseudo galvanic-HBC behaviour owing to the presence of a pair of electrodes attached to the body. The behaviour causes the HBC source to drop only a part of its potential, *V*_*HBC*_, across *C*_*L*_, the rest drops across *R*_*EH*_. The voltage across *C*_*L*_ divides between *C*_*G*_ and *C*_*B*_. The voltage across the *R*_*EH*_ divides between 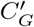 and *C*_*B*_. The net voltage across *C*_*B*_ is the sum of the voltage drops across *C*_*B*_ contributed by voltages across *C*_*L*_ and *R*_*EH*_. Equations 2, 3, and 4 describe the voltage across *C*_*L*_, *R*_*EH*_, and *C*_*B*_. Equation 5 represents the transmitter-side channel gain, i.e., *V*_*b*_*/V*_*HBC*_.

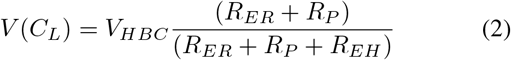

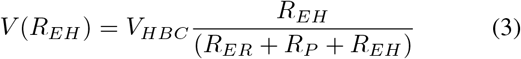

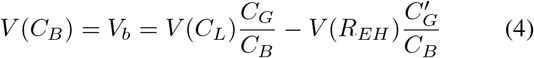

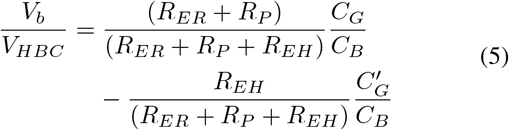

Figure 5 (C) plots the variation in the transmitter-side channel gain with *R*_*P*_, *R*_*ER*_, and *R*_*EH*_. We find that the presence of RLD (*R*_*p*_ < *∞*) affects the channel gain. We find increased channel loss with lower *R*_*p*_. We also note that channel loss increases with an increase in *R*_*ER*_ or *R*_*EH*_.

#### 2) Experiments

Our theory predicts that RLD increases channel loss. This section conducts a series of experiments to analyze this effect. The experimental setup consists of a floating on-body node with an ECG sensing frontend (supporting RLD) and an HBC transmitter. The node transmits HBC signals for different RLD configurations. For each of these configurations, we use an on-body HBC receiver to measure the strength of the HBC signal at two different locations on the body: chest and right palm. The experiments only observe the effect of the RLD on HBC and do not record meaningful ECG. Figure 6 summarizes the different experiments performed and their results. In the subsequent paragraphs, we describe these experiments and our findings.

**Fig. 6.**
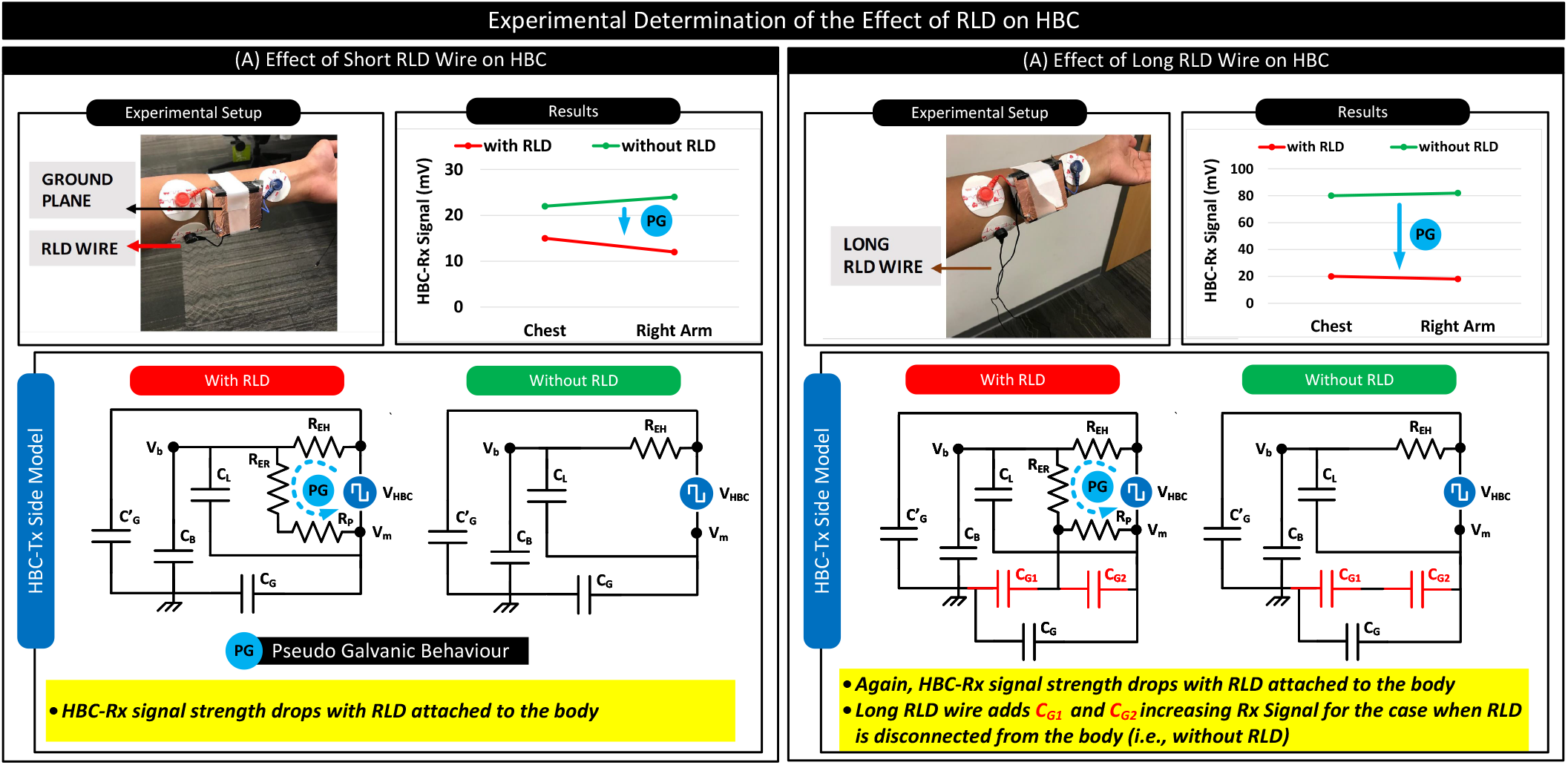
Experimental setup, circuit model and results to analyse the effect of RLD on HBC. In the experiments, the HBC transmitter is placed on the left arm, and the receiver is placed on the chest and right arm. The received signal strength is noted for cases with RLD connected to the body and for the case when it is disconnected from the body for different RLD wire lengths. We note that the received signal strength from both the chest and right arm is lower when RLD is attached to the body than when it was left floating due to the pseudo galvanic behaviour.

##### a) Experiments with Short RLD Wire

In this experiment, a short RLD wire is connected and disconnected from the body to see the variations in the received HBC signal strength. Figure 6 (A) shows the experimental setup, the circuit models and the received signal strength. When RLD is attached to the body, it creates a low-resistance return path for the transmitted HBC signals, mimicking a pseudo galvanic behaviour. The behaviour results in only a part of *V*_*HBC*_ dropping across *C*_*B*_ resulting in higher channel loss and a lower received signal. We find that the experimental results align with the theory. We note that the received signal strength from both the chest and right arm is lower when RLD is attached to the body than when it was left floating.

##### b) Experiments with Long RLD Wire

In this experiment, we replace the small RLD wire with a long RLD wire. Again, we note that RLD deteriorates signal strength, similar to the results from the previous experiment. We note that when the RLD wire is left floating, the signal strength is higher than the previous case, which uses a short RLD wire. We infer that a floating RLD wire increases the ground plane surface area, adding capacitances, *C*_*G*1_ and *C*_*G*2_ across *C*_*G*_ resulting in higher channel gain and received signal strength.

### Conclusion

Our findings from the theory and experiments with RLD are as follows. For a node performing simultaneous sensing and communication, RLD affects HBC. RLD creates a low resistance return path for the transmitted HBC signals, mimicking a pseudo-galvanic-HBC behaviour resulting in higher channel losses for capacitive-HBC irrespective of the length of the RLD wire.

## IV. RLD-Free Sensing — Justification

In Section III, we show that RLD affects HBC transmissions. Notably, the presence of the RLD electrode on the body increases channel loss. We propose not to use RLD for biopotential sensing in wearables as a solution. We argue that due to wearables’ floating nature, the effect of powerline noise on biopotential measurements is drastically lower than in the case of grounded measurement systems such as the medical-grade device shown in Figure 4 (A). Further, we propose using software filters such as the notch filter to remove any residual 50 Hz/60 Hz noise in the absence of RLD. We expect our solution to be practical since nodes in a BAN have a wearable form factor (with floating grounds) in most applications. In this section, we put forward the theory reasoning why RLD is unnecessary for a wearable performing biopotential sensing.

RLD, as described in Section II-B, is used for two purposes. To fix the ECG signals’ common-mode and attenuate common-mode variations due to power line interferences with the body, affecting ECG measurements. A possible way to fix common-mode without RLD is to use AC coupling with DC bias resistors. Alternately, one may also rely on the contact potential of the electrodes. Concerning attenuating power line interference, ECG demand a 92 dB Common Mode Rejection (CMR) [23]. The ECG sensing systems achieve the required CMR of 92 dB in two parts. The parasitic pathways that couples the power line potential to the ECG sensing system and RLD contribute to one part of the CMR. Depending on its CMRR specification, the difference amplifier (with its passives) adds the other part of the CMR.

Figure 7 (A) shows the generic CMR model for an ECG sensing system [28]. The CMR of the sensing system is *−*20*log*_10_(*V*_*out*_*/V*_*in*_). Where *V*_*out*_ is the output voltage of the difference amplifier and *V*_*in*_ = 110 V, being the powerline potential. Here, CMR comprises two components: *CMR*_1_ and *CMR*_2_. *CMR*_1_ is the common-mode rejection added by the parasitic pathways that couple the powerline signal to the ECG sensing system and the RLD. *CMR*_2_ is the common-mode rejection achieved by the difference amplifier depending on its Common-Mode Rejection Ratio (CMRR) specification. For a typical CMRR of 60 dB (effective CMRR of the difference amplifier with passives) and a gain setting of 20 dB, the amplifier results in a common-mode rejection, *CMR*_2_ of 40 dB [23]. Since 92 dB is the target CMR to achieve, for a *CMR*_2_ of 40 dB, the required *CMR*_1_ is 52 dB. Figure 7 (B),(C) derives the *CMR*_1_ for a grounded and a wearable (i.e., floating) ECG sensing system for typical parasitic settings and RLD protection resistor of 2 MΩ. We note that the presence of RLD with and without a feedback loop improves *CMR*_1_ for both grounded and wearable systems. We find that owing to the floating nature of the wearable system (i.e., *C*_*G*_*≠ ∞*), *CMR*_1_ of the wearable system is higher than *CMR*_1_ of the grounded system for similar settings. We find that *CMR*_1_ of the wearable system without RLD is 77 dB, which is more than *CMR*_1_ of 52 dB required for an ECG sensing system. We find that *CMR*_1_ of the wearable system without RLD closely matches *CMR*_1_ of the grounded system with RLD and feedback loop.

**Fig. 7.**
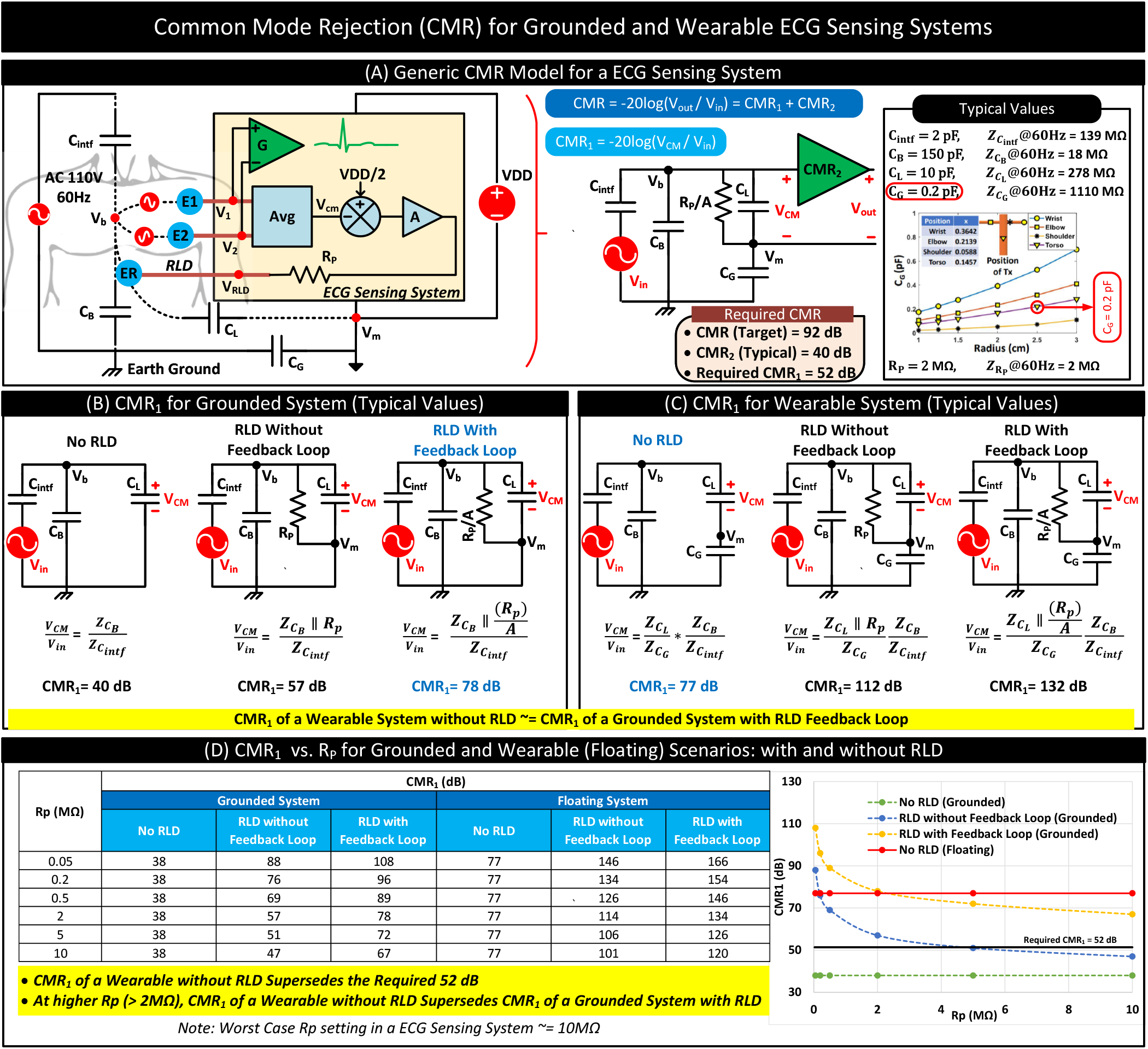
(A) Generic CMR model of a ECG sensing system comprising of *CMR*_1_ (contributed by parasitic pathways and RLD) and *CMR*_2_ (contributed by the difference amplifier and its passives), with typical values for the models parasitic components [27]. The graph C_G_ vs Radius is reprinted from Datta, Arunashish, et al. “Advanced biophysical model to capture channel variability for eqs capacitive hbc.” IEEE Transactions on Biomedical Engineering 68.11 (2021): 3435-3446 [27]. (B) *CMR*_1_ for grounded and wearable systems with and without RLD. We note that *CMR*_1_ for a wearable system without RLD closely matches the *CMR*_1_ of a grounded system with RLD and feedback loop.(C) Variation in *CMR*_1_ with RLD protection resistor *R*_*p*_. At higher *R*_*p*_ (2 MΩ), *CMR*_1_ of a wearable without RLD Supersedes *CMR*_1_ of a grounded System with RLD and feedback loop.

Figure 7 (D) shows the variation in *CMR*_1_ for different values of *R*_*p*_. At higher *R*_*p*_ (2 MΩ), *CMR*_1_ of a wearable without RLD Supersedes *CMR*_1_ of a grounded System with RLD and feedback loop. Since the worst-case Rp can be as large as (10 MΩ) [29], the wearable system without RLD has a superior *CMR*_1_ performance compared to a grounded system with RLD for most practical settings.

In summary, although RLD can improve the *CMR*_1_ of a wearable ECG sensing system, it is not a requirement. Since the *CMR*_1_ of a wearable system without RLD supersedes the *CMR*_1_ requirement of an ECG sensing system and the *CMR*_1_ of grounded systems with RLD and feedback loop.

## V. Proof-of-Concept Wearable Design: RLD-Free ECG Sensing Supporting HBC

In Section III, we show that RLD affects HBC transmissions, increasing channel loss. As a solution, we propose not to use RLD for wearables supporting HBC. We backup our proposed solution in Section IV by arguing that RLD is not a requirement for wearable ECG sensing systems with a floating ground common in BAN settings. In this section, we validate our argument. For the validation, first, we develop a proof-of-concept wearable that measures ECG without RLD. The wearable uses filters to substitute the RLD functionality to reduce interference. Next, we conduct experiments to verify the wearable’s quality of ECG sensing and HBC transmissions. We demonstrate that with appropriate filters in place, in wearables, RLD-free sensing does not impact the quality of the ECG signals. Further, HBC can transmit ECG signals with a 96% correlation compared to BLE at a fraction of the power.

### A. Wearable Design

Figure 8 shows the components of our wearable sub-inch^3^ custom-designed node. The node is a modified version of the wearable described in our previous work [11]. It can carry single-lead ECG measurements without RLD and transmit the ECG samples via HBC and BLE in real-time. The node uses BLE as the gold standard to compare HBC transmissions.

**Fig. 8.**
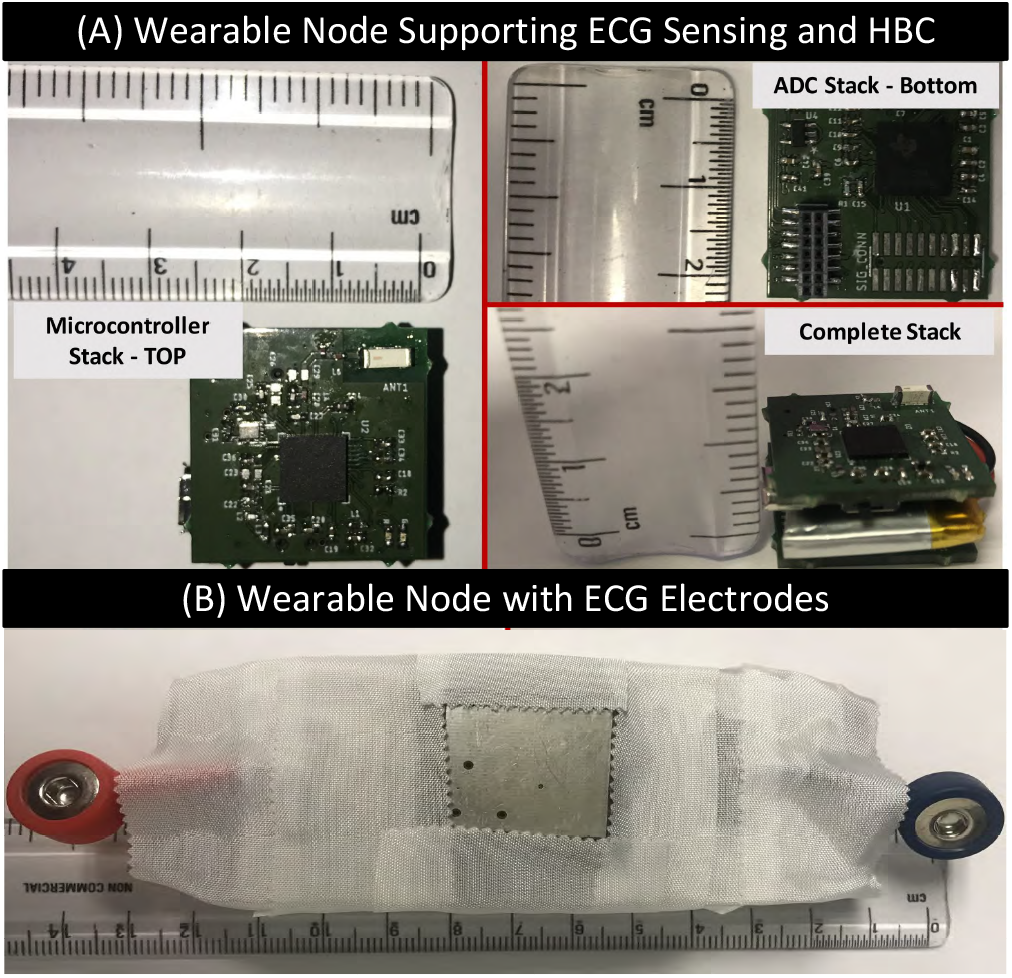
Device Design: a) Custom designed sub-inch^3^ wearable node capable of sensing and transmission of ECG signal using HBC and BLE. b) Complete device setup with ECG electrodes.

The node consists of two vertically stacked PCBs. The bottom PCB consists of IC ADS1298 from Texas Instruments. The IC ADC1298 includes a front-end differential amplifier to acquire a single-lead ECG from the body and a 24-bit ADC to digitize the signals. For ECG sensing, two Ag-AgCl hydrogel electrodes, attached to the chest and separated by a distance, are connected to the inputs of the difference amplifier of ADS1298 via two snap connectors on the PCB. Figure 8 shows these snap connectors; the RED and BLUE snap connectors interface the electrodes attached to the right and left regions of the chest, respectively. The node uses RC low pass filters at the input terminals of the difference amplifier to eliminate high-frequency noise, including HBC from ECG. To couple HBC signals to the body, the bottom PCB uses a signal plane (a copper pad) located between the snap connectors.

The top PCB consists of the SoC NRF52833 with an inbuilt BLE radio from Nordic Semiconductors. The SoC periodically reads the 24bit ECG samples from ADS1298 via SPI and transmits them over BLE and HBC. NRF52833 transmits ECG samples via BLE as a string of characters with a newline character to separate samples. NRF52833 transmits ECG samples via HBC in binary OOK format. A 500 kHz, 50% duty cycle square wave was used as the carrier. The node achieved an effective HBC transmissions rate of 25 Kbps.

### B. Experiments

To validate our wearable node’s ECG sensing and HBC transmission capabilities, we perform two experiments. First, to demonstrate the quality of ECG measurements without RLD, and second, to validate the quality of HBC transmissions by comparing the ECG data sent via HBC with BLE. For the experiments, we use the setup in Figure 9(a). It consists of a subject wearing the node with the ECG electrode and HBC transmitting electrode in contact with the chest. The node periodically measures ECG samples and transmits them simultaneously over HBC and BLE. We use A BLE receiver connected to the computer for receiving transmitted BLE data. We use a computer-based oscilloscope by Pico Technologies to probe the subject’s wrist at 3.9 MSamples*/*s to receive the transmitted HBC signals in OOK format. We then post-process these signals in Matlab to decode the transmitted HBC data. First, the sample points from the oscilloscope are bandpass filtered between 400 KHz-to-600 KHz with 80 dB attenuation. Next, we use envelope detection and thresholding techniques to decode the signals from the filtered sample points. Additionally, we also use software error-correcting algorithms to minimize possible errors while decoding HBC data.

**Fig. 9.**
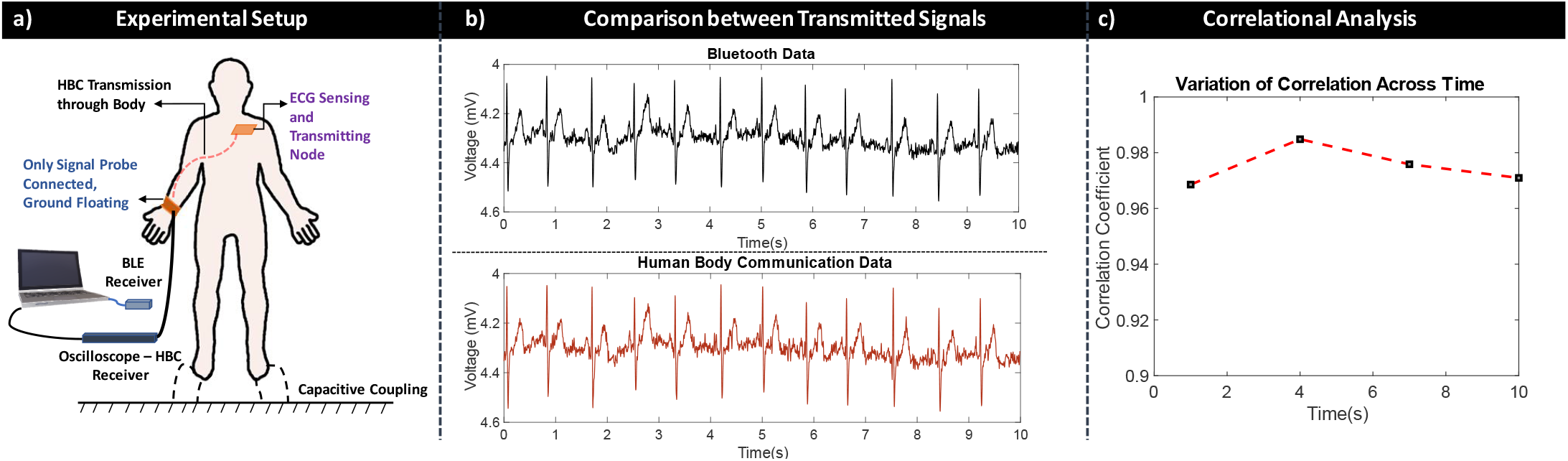
Experimental Results: a) Experimental Setup depicting the setup used to transmit biopotential signals using Bluetooth and HBC. b) ECG results for the Bluetooth transmitted data and HBC transmitted data. c) Correlation coefficients between Bluetooth and HBC

#### 1) RLD-Free ECG Sensing

We note that without RLD, ECG signals are affected by noise. We, however, find that a combination of proper hardware and software filtering techniques as in Figure 10 can drastically reduce the noise. We describe these filters and their results below.

**Fig. 10.**
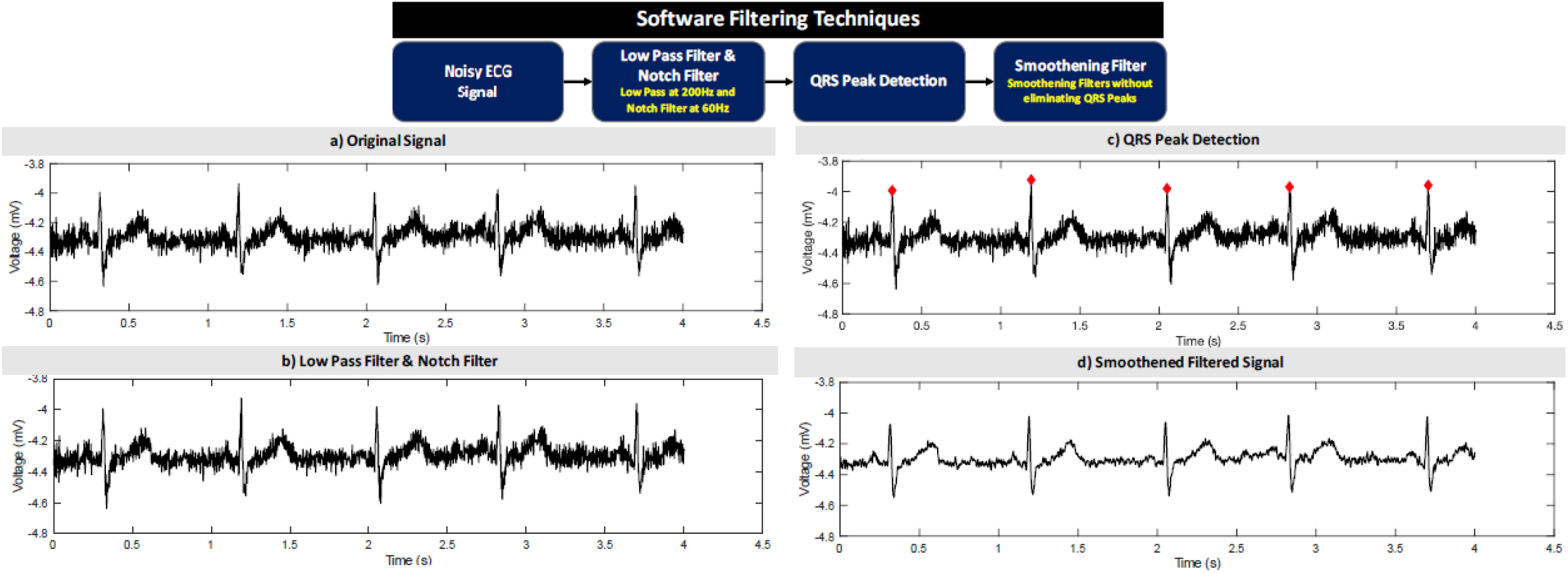
Software filtering to remove noise: a) Original unfiltered signal (Noisy ECG signal), b) Low Pass Filter and Notch filtering to eliminate 60Hz noise, c) QRS peak detection to isolate peaks to prevent over filtering, d) Smoothening techniques to eliminate noise in the signal.

#### Hardware Filter

We use the first-order RC filters at the input terminal of the differential amplifier of ADS1298 to remove high-frequency noise sources such as HBC received by the ECG electrodes from corrupting ECG measurements. We configured these filters for a cut-off frequency of 230 Hz. *Software Filter:* We use a software notch filter to eliminate 50 Hz/60 Hz line interference. Further, QRS complexes in the ECG signal are detected, and smoothening techniques are employed to eliminate spurious spikes in the ECG signal. In this technique, to prevent the filters from considering the QRS peaks as spurious spikes, the markers for the QRS complexes are detected first. We then bypass the sample points corresponding to these from filtering. Figure 10 shows the ECG signal at different stages of the software filtering.

### 2) HBC — Quality of Transmissions

For medical-grade setups, wired communication technologies generally have the best SNR; they are often used to benchmark new communication technologies for medical applications. However, using wired communication technologies to benchmark HBC is problematic as the wires can act as grounds and vary the HBC channel characteristics. As a solution, in this work, we evaluate the quality of HBC by comparing it with BLE, generally considered a gold standard communication modality for biopotential signal transfers. Figure 9 shows the comparison of ECG signals from HBC and BLE. Results indicate a high correlation of > 96 %, indicating the suitability of HBC for biomedical applications.

## IV. Conclusion

For the first time, our work studies the interaction between Human Body Communication (HBC) and biopotential sensing systems that uses Right Leg Drive (RLD). We find that HBC, owing to its higher frequency content, does not affect the sensing of biopotential signals from the body. The anti-aliasing low pass filter at the inputs of the sensing circuit blocks HBC signals avoiding possible interactions HBC can have with biopotential signals. We note that when a node performs biopotential sensing using Right-Leg Drive (RLD), it affects its HBC transmissions. RLD electrode, in this case, offers a low impedance return path for the HBC signals increasing channel loss. We suggest RLD-free sensing of bipotential signals in wearables supporting HBC as a solution. Due to their floating nature, we show that wearables are less prone to powerline noise interferences and hence do not require RLD. We validate our proposed solution using a proof-of-concept wearable which performs ECG sensing without RLD.

## VII. Acknowledgment

This research is supported by the National Science Foundation Career Award under Grant CCSS 1944602. The experimental studies on humans are approved by the Purdue Institutional Review Board (IRB) #1610018370.

## Notes

### Competing Interest Statement

The authors have declared no competing interest.

### Summary of Updates

Updated Figure-7 caption with proper citation to references.

